# Causally Linking Neural Dominance to Perceptual Dominance in a Multisensory Conflict

**DOI:** 10.1101/2020.04.06.027227

**Authors:** Kyongsik Yun, Joydeep Bhattacharya, Simone Sandkuhler, Yong-Jun Lin, Sunao Iwaki, Shinsuke Shimojo

## Abstract

When different senses are in conflict, one sense may dominate the perception of other sense, but it is not known whether the sensory cortex associated with the dominant modality exerts directional influence, at the functional brain level, over the sensory cortex associated with the dominated modality; in short, the link between sensory dominance and neuronal dominance is not established. In a task involving audio-visual conflict, using magnetoencephalography recordings in humans, we first demonstrated that the neuronal dominance – visual cortex being functionally influenced by the auditory cortex – was associated with the sensory dominance – participants’ visual perception being qualitatively altered by sound. Further, we found that prestimulus auditory-to-visual connectivity could predict the perceptual outcome on a trial-by-trial basis. Subsequently, we performed an effective connectivity-guided neurofeedback electroencephalography experiment and showed that participants who were briefly trained to increase the neuronal dominance from auditory to visual cortex also showed higher sensory, i.e. auditory, dominance during the conflict task immediately after the training. The results shed new light into the interactive neuronal nature of multisensory integration and open up exciting opportunities by enhancing or suppressing targeted mental functions subserved by effective connectivity.

## Introduction

We continuously encounter with visual and auditory information, processed by distinct sensory cortices, which are eventually integrated to produce a conscious behavioral unique response [1, 2]. However, when visual and auditory information is incongruent or in conflict, one sensory modality may dominate the other, leading towards a multisensory illusion [3]. A critical question remains whether sensory dominance is linked to neuronal causality, i.e. sensory cortex of the dominant modality would causally influence, at the functional level, the activities of the sensory cortex of the subordinate modality.

We tested this specific prediction in the framework of an audio-visual conflict – sound-induced flash illusion [4, 5]: a multisensory illusion, when a single flash in the visual periphery is accompanied by two beeps, the single flash is often misperceived as two flashes. Individual differences in proneness to the illusion are reflected in the neurochemical [6] (GABA concentration in superior temporal sulcus), structural [7] (grey matter volume in early visual cortex), and functional excitability [8, 9] (visual event-related responses to sound) differences. However, these findings do not explain the trial-by-trial variability, i.e. observers perceive the illusion sometimes, but not always, even though the physical stimuli remain identical and supra-threshold across trials. Since the auditory information dominates over the visual information for this illusion to occur, neural activity in the auditory cortex is predicted to exert a causal influence on the activity in the visual cortex, not the other way around.

We addressed this question by recording MEG signals from healthy humans in the sound-induced flash paradigm (Fig. 1A). We compared the effective connectivity between auditory to visual cortices for illusion and non-illusion trials, differing only in terms of the qualitative nature of visual perception, and verified our prediction. Next, to establish a causal mechanism, we performed a separate experiment involving EEG based neurofeedback in which participants were briefly trained to spontaneously regulate their auditory to visual effective connectivity and found that such connectivity-based neurofeedback training significantly increased the probability of auditory stimulus qualitatively altering the visual perception. MEG was used to quantify the trial-by-trial effective connectivity between auditory and visual cortices due to its high sensitivity, and EEG was used as a neurofeedback tool to modulate the effective connectivity due to its practicality.

**Figure 1.**
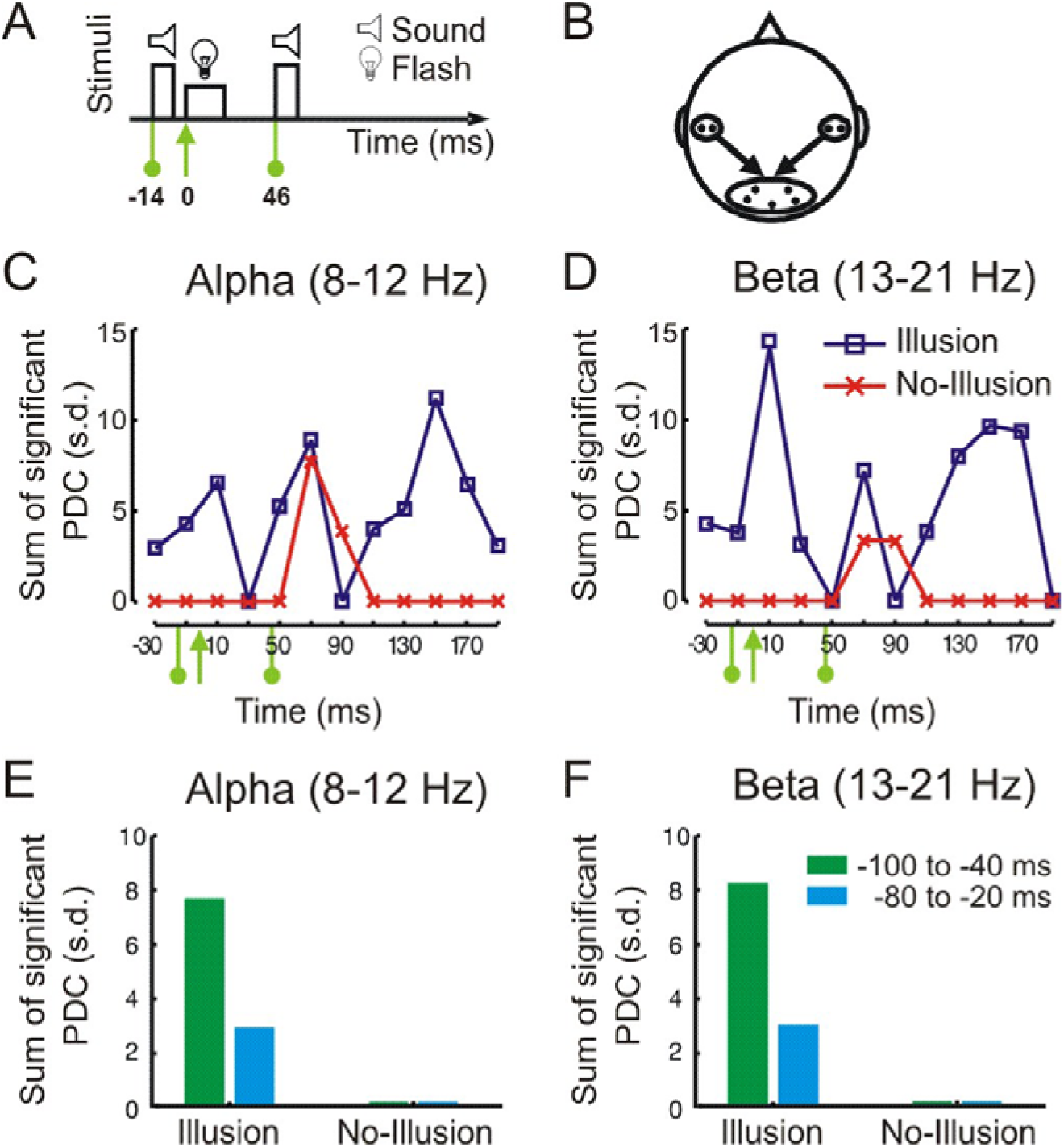
Experimental setting of sound-induced flash illusion, and strong partial directed coherence from auditory to the visual cortex, but primarily in illusion trials. (A) Sound-induced flash illusion stimuli parameters. The auditory stimulus consisted of two brief beeps each lasting 10 ms and separated by 50 ms. The flashing stimulus was a uniform white disk appearing in the periphery (8.5° eccentricity) for a duration of 20 ms. (B) Considered sensors and direction of information flow. (C)-(D) Sum of significant PDC values (rank test; *p* < 0.005, see Experimental procedures), expressed in s.d., displaying the degree of the causal influence of auditory cortex onto visual cortex in (C) alpha (8-12 Hz) and (D) beta band (13-21 Hz) as a function of time. Each time point corresponds to a time-window spanning ± 30 ms. For example, the first time-point at −30 ms spans a time-window from −60 to 0 ms with respect to flash onset. Green markers indicate flash and auditory beep onsets (see (A)). (E)-(F) Sum of significant PDC values (rank test; *p* < 0.005) from auditory cortex to the visual cortex in the −100 to −40 ms and −80 to −20 ms pre-flash-onset time window in (E) alpha and (F) beta band.

## Materials and methods

### Ethics statement

All participants provided written informed consent before the experiments and were paid for their participation. The MEG study was approved by the Internal Review Board of National Institute of Advanced Industrial Science and Technology, Osaka, Japan, and the EEG study were approved by the Internal Review Board at California Institute of Technology, Pasadena, USA; both studies were conducted following the Declaration of Helsinki.

### Participants

For the MEG study, 11 adults (3 females, ages ranging between 22-40 years) participated. For the EEG study, 27 adults (11 females, ages ranging between 22-40 years) participated. The sample sizes were comparable to previously published related studies [5, 10]. Two sets of participants were completely independent. All participants were healthy, had no history of neurological or psychiatric disorders and had normal or corrected to normal visual acuity, and normal hearing.

### MEG study: Design, procedure, and materials

The MEG signals were recorded with a 122-channel whole-scalp planar-gradiometer (Neuromag 122, Elekta-Neuromag Oy, Helsinki, Finland) in a magnetically shielded room. The instrument measured two orthogonal tangential derivatives of the magnetic field at 61 scalp locations. In the examined bimodal condition, the event trigger was synchronized with the onset of the flash. The subjects were seated upright with their heads comfortably resting against the inner wall of the helmet and were instructed to fixate on a cross on the screen, and not to blink during trials.

The experiment consisted of four conditions: (i) a visual flash, (ii) a flash accompanied by two auditory beeps, (iii) two beeps and no flashes, and (iv) two flashes. The flashing stimulus was a uniform white disk subtending a visual angle of 2° in the periphery at 8.5° eccentricity for a duration of 20 ms. The auditory stimulus consisted of two brief beeps each lasting 10 ms and separated by 50 ms. The sound stimulus (1 kHz frequency at 70 dB SPL) was presented by headphones. In the bimodal condition, the flash onset was 14 ms after the onset of the first beep. There were 80 trials for each condition and the order of the trials was random. The inter-trial interval was varied randomly between 1500 and 2000 ms. The participant’s task was to judge the number of flashes they perceived at the end of each trial in a three-response-category paradigm – zero, one, or two flashes.

The continuous MEG signals were band-pass filtered at 0.01 – 100 Hz, digitized at 550 Hz and stored for off-line analysis. To remove the contamination due to spurious oscillations (∼ 40 Hz) of Helium cylinders, a further band-pass filtered was applied at 0.05 – 30 Hz using a Butterworth filter of order 3. The epochs containing eye blinks or excessive movements were excluded based on amplitude criteria. Here, we considered only one experimental condition, a flash accompanied by two beeps that have two possible outcomes: (i) no-illusion – perceiving one flash, (ii) illusion: perceiving two flashes.

We used partial directed coherence, PDC [11] to identify the direction of information flow. Multivariate autoregressive models were adaptively estimated using overlapped time-windows (60 ms time-windows with 40 ms overlap) to make the estimated model parameters varying smoothly. The optimal model order was determined by locating the minimum of the Akaike Information Criterion (AIC) [12] across time and was set to 6. Statistical significance of PDC values was determined by independently shuffling the trial order across participants for each sensor. Thus, we obtained PDC values that were due to chance by pooling over participants. The data were shuffled for 200 times, and we used a nonparametric rank test as a qualitative measure of significance. Only for those PDC values that passed this nonparametric test, we expressed significant PDC values in terms of standard deviations of the shuffled distribution to have better visual clarity of the degree of causal interdependence.

For predicting the perception of one (*θ* = 1, i.e. no-illusion) or two flashes (*θ* = 2, i.e. illusion), we applied a Bayesian classifier with a uniform prior probability. Input data for this classifier was the directed influence from AC (4 sensors) to VC (5 sensors) (see Figure 1B). For predicting perceptual outcome on a trial-by-trial basis, we estimated PDC on each trial. Here, we considered bivariate autoregressive models (with optimal AIC model order of 3) and longer (i.e. 100 ms) time-windows to get reliable estimates. The immediate pre-stimulus time-window was −114 ms to −14 ms and the post-stimulus time-window was 0 to 100 ms.

The random variable *y* represents the classification input data vector of PDC values in alpha and beta bands. Bayes’ Theorem gives us the posterior probability of *θ* given the information that *y* occurred:

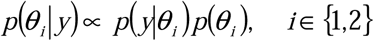

where *p*(*θ*_*i*_) is the prior probability of *θ*_*i*_, which is uniform by design and *p*(*y*|*θ*_*i*_) is the probability distribution of *y*, which we estimated by a Gaussian mixture model with two components. The predicted post-stimulus response was subsequently chosen to be the one with maximum probability. We repeated 10-fold cross-validation 100 times to assess the performance of the classification accuracy.

### EEG study: Design, procedure, and materials

Each participant was seated in front of the computer screen. The EGI (Electrical Geodesics Inc., Eugene, OR) cap was used for the EEG recording and analysis. The experiment consists of three sessions: pre-training, neurofeedback training, and post-training sessions. First, in the pre-training session, participants were instructed to answer using a keypad how many flashes they perceived and they performed 100 trials of sound-induced visual illusion tasks. In the center of a 15-inch black computer screen, 20×20 mm sized white crosshair (+) was shown across all the trials and participants were asked to look at the crosshair during all the tasks. On each trial, a 67 mm diameter white circle appeared at the bottom of the screen for 16 ms. The first beep was played 14 ms before the white circle appeared. Then the second beep was randomly played 46 ms after the white circle appeared. Inter-trial interval randomly varied between 1 s to 3 s.

Next, participants were randomly assigned to one of the two groups: A→V and V→A training groups. Participants of A→V training group were shown a bar graph displaying the real-time processed A→V connectivity of their brains. They were asked to try to figure out how to increase the height of the bar graph. Participants of V→A group was shown the bar graph displaying V→A connectivity. In essence, the participants were only instructed to “control” their brain connectivity voluntarily and heighten the bar graph on the computer screen. The neurofeedback training lasted for a brief period of 5 min. Subsequently, participants performed the post-training tasks that were the same as they did before the EEG neurofeedback training.

EEG was recorded at a sampling rate of 1000 Hz using 128-channels EGI cap. The EEG activities at 7 channels (T3, T4, T5, T6, O1, O2, and Oz) between 8-12 Hz were used for PDC computation. The impedance of the electrodes was kept below 50 kΩ. Real-time frequency filtering to extract alpha frequency band (8-12 Hz) and the PDC computation were performed. The processing latency was 223ms +-26ms. The detected EEG signal was both recorded for analysis and fed back to the subject forming a feedback loop. Computed connectivity using PDC from auditory (T3, T4, T5, T6) to visual cortices (O1, O2, Oz) was represented as the height of the bar graph and its sign was reversed at the bar graph shown to the control group. While participants tried to heighten the bar graph, their brain connectivity was modulated and in turn, formed the feedback loop.

## Results

### Experiment 1: MEG study linking neural dominance to perceptual dominance Auditory to visual connectivity was associated with the double-flash illusion

Flash illusion was reported for 62% of trials (i.e. out of 687 trials, participants reported perceiving two flashes on 424 trials), while stimulus parameters remained identical with 2 beeps and 1 flash (Fig 1A). We used partial directed coherence [11], a frequency domain representation of Granger’s causality [13], to measure the effective connectivity (i.e. the explicit and directional flow of information) between auditory and visual cortices. We focused our analysis in the alpha (8-12 Hz) and the beta (13-21 Hz) band neuronal oscillations after previous studies [10, 14]. With the adaptive multivariate autoregressive modeling approach for short window spectral analysis [12], we determined the connectivity from the nine selected MEG sensors located approximately over the auditory cortex (AC) and visual cortex (VC) (Fig 1B). We observed a robust flow of information from auditory to the visual cortex for the illusion trials in both alpha (Fig 1C) and beta (Fig 1D) oscillations; on the other hand, such directional flow of information from auditory to visual cortex remained mostly non-significant (except around 70 ms after flash-onset). The timings of the peaks of auditory to visual connectivity at 40 to 100 ms [15, 16] and 110 to 170 ms [15] for illusion trials are in close agreement with the reported time-intervals of previous studies on multisensory integration. However, in contrast to earlier findings [15, 16] which compared multisensory to unisensory conditions, we compared two identical multisensory conditions, differing only in the quality of the subjective perception. Therefore, our results establish a direct link between the brain’s specific connectivity pattern and conscious awareness. This potentially causal influence on the visual cortex by the auditory cortex at such an early stage of information processing may be indicative of direct communication between these two sensory areas at a functional level. Of note, earlier studies [17, 18] suggest direct structural connectivity between these two sensory areas, especially between the primary cortices. Both studies reported that these projections target the peripheral visual field representation in the visual cortex, which matches with our earlier results [4] that the sound-induced flash illusion is stronger if the visual flash is presented in the periphery than in the fovea.

### Directedness and asymmetrical nature of auditory to visual connectivity

To validate that these causal functional modulations were possibly direct and not via other multisensory areas, we repeated the connectivity analysis after including different sensors from other multisensory regions including parietal, frontal, and temporal cortex in our information flow model (see Figure 2A-B; left panel) and by omitting some sensors from AC and VC areas. Results for different model configurations are shown in Figs. 2(A-B) and Figs. 2(C-D) for alpha and beta band, respectively. Despite the variations in the temporal profiles from AC to VC connectivity across model configurations, we observed that overall the degree of AC to VC was larger and more sustained in the illusion trials than no-illusion trials, thereby confirming our earlier findings. Thus, the reported early AC to VC connectivity was unlikely to be influenced by the higher-order multisensory areas.

**Figure 2.**
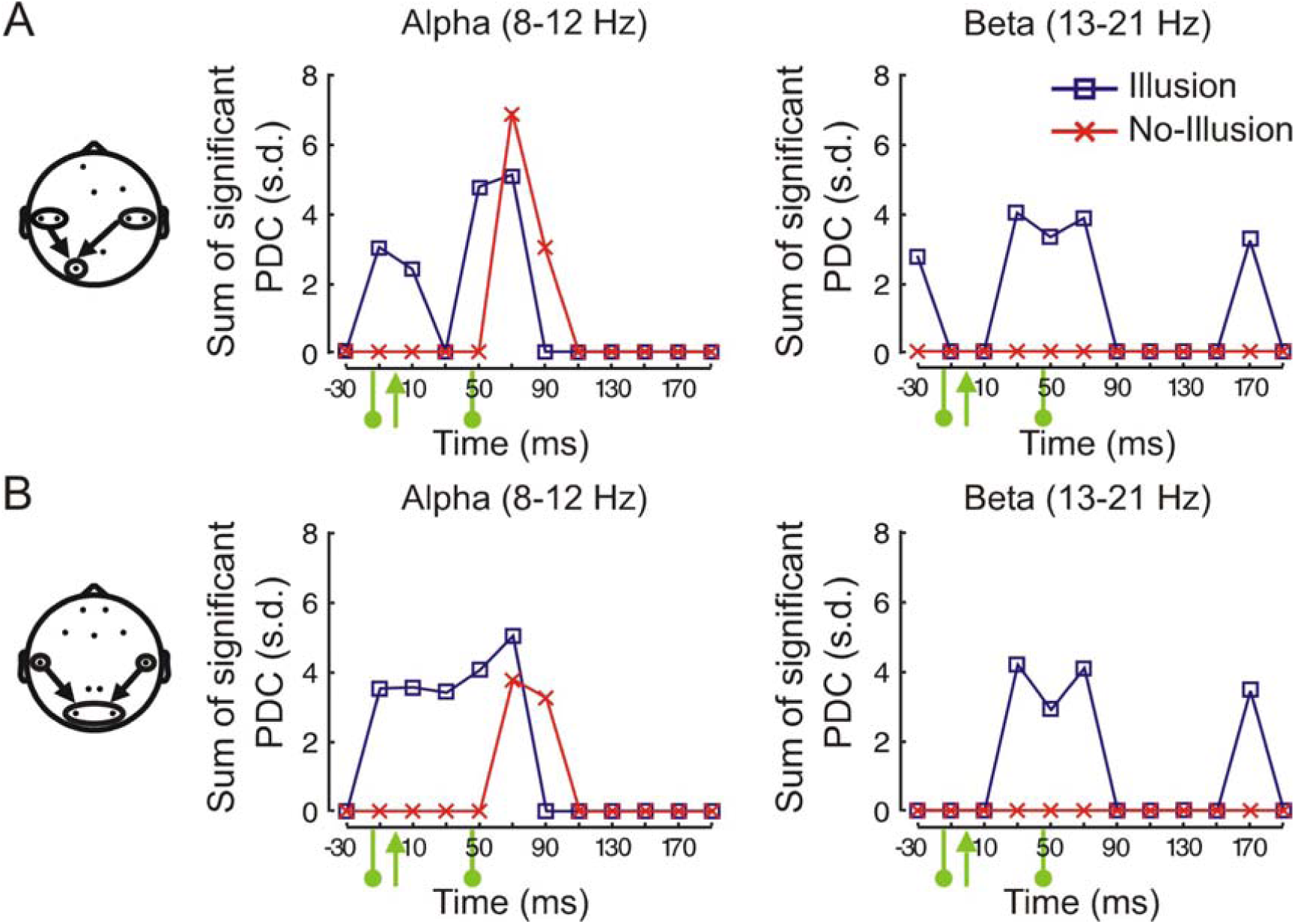
Two control sensor settings to investigate potentially directed nature of the influence from AC to VC. (A) Left, considered sensors and direction of information flow. Some AC and/or VC sensors were omitted for both settings to constrain the dimension of the multivariate AR model. Sensors that showed the strongest responses in ERP analysis were included. Right, the sum of significant (rank test; *p* < 0.01) PDC values, expressed in s.d., display degree of the causal influence of AC onto VC in alpha (8-12 Hz) and beta band (13-21 Hz) as a function over time (see Figure 1C-D). (B) As in (A) for second sensor setting incorporating bilateral sensors.

Next, we inspected the connectivity in the reverse direction, i.e., the influence of the visual cortex onto the auditory cortex. In the flash illusion, sound dominates vision, but not vice versa. Aligned with this inherent nature of the illusion, we found that the information flow from the visual cortex to the auditory cortex was comparable between illusion and non-illusion trials (see Figure S1, Supplemental Digital Content). This suggests that the effective connectivity from AC to VS, but not the other way round, is crucial to alter the qualitative nature of visual perception in the sound-induced flash illusion.

### Prestimulus auditory to visual connectivity predicting perceptual outcomes

Given the early nature of the causal interactions, and the recently reported evidence of pre-stimulus brain states shaping post-stimulus responses [19-21], we investigated the immediate pre-stimulus period (100 ms before flash-onset) and found robust differences between illusion and non-illusion trials (Figure 1C, D). In illusion trials only, we found strong causal influence exerted by the auditory cortex onto the visual cortex in the pre-stimulus period. We suggest, therefore, that the spontaneous fluctuations of this causal interaction between two sensory cortices in the prestimulus period might bias sensory perception in ambiguous or sensory-conflicting situations

If the effective connectivity from auditory to visual cortex has a causal role in biasing decisions, it would be possible to predict, above chance, the behavioral response from the connectivity values on a trial-by-trial basis. We tested this by applying a machine-learning technique. Using PDC values in the alpha and beta frequency bands (estimated from 100 ms long time-windows) as features in a Bayesian classifier, we predicted the behavioral response (either illusion or no-illusion). Using the pre-sound onset time window only gave an accuracy of 55.3 % (one-sided exact binomial test, *n* = 68700, successes = 37998, H_0_: probability of success = .5; *p* < 0.0001), whereas using the immediate post-flash onset time-window decreased (Mann-Whitney, *p* < 0.0001 with respect to pre-stimulus time-window) accuracy to 53 % (successes = 36247, *p* < 0.0001). However, when using the joint information from that pre- and post-stimulus onset time-window, the mean prediction accuracy improved to 61.4 % (successes = 42184, *p* < 0.0001). Although this classification accuracy is relatively moderate (possibly due to our simple model excluding brain regions other than AC and VC, a brief period, and less robust estimation of PDC values at the single-trial level), the prediction improvement, after including the immediate pre-stimulus period, remained statistically significant.

These results, altogether, provide robust and consistent evidence that the effective connectivity from the auditory to the visual cortex significantly induces a qualitative alteration of visual perception by sound in the sound-induced flash illusion.

### Experiment 2: EEG based effective connectivity guided neurofeedback causally modulating perceptual dominance

To establish a piece of further causal evidence for this link between neural dominance and perceptual dominance, we subsequently performed an effective connectivity-guided neurofeedback EEG experiment (*n*=27) consisting of three sessions: pre-training, training, and post-training. In the pre-training session, participants were presented with 100 trials each of the four conditions: 1 flash with 1-4 beeps; participants had to report the number of perceived flashes on each trial. In the brief training session (5 min [22]), the participants were shown a bar graph displaying the real-time effective connectivity measure, either auditory to the visual cortex, A→V, or visual to the auditory cortex, V→A, as measured by PDC in the alpha band. The participants were instructed to increase the height of the bar graph by voluntarily “controlling” the level of spontaneous audio-visual alpha band cortical connectivity. The EEG activities at 7 electrode locations (auditory: T3/4, T5/6; visual: O1/2, Oz) were used for PDC calculation in the alpha band (8-12 Hz) after previous studies [14] and our MEG findings. Half of the participants increased A→V cortical connectivity and the other half increased V→A connectivity. The post-training session was immediately after the training sessions, and the participants were presented with the same task as in the pre-training session.

Next, we investigated whether this information flow indeed occurred during the sound-induced flash illusion and whether information flow changes after connectivity-based neurofeedback training. The PDC of A→V connectivity in illusion trials was significantly larger than in non-illusion trials (*t*(26)=2.21, *p*=0.036), while PDC of V→A connectivity did not differ significantly between illusion and non-illusion trials (*t*(26)=0.062, *p*=0.95) (Figs. 3C,D). So, our earlier MEG findings of linking neural dominance, from auditory to the visual cortex, to perceptual dominance, sound modulating vision, was replicated using EEG from an independent sample.

**Figure 3.**
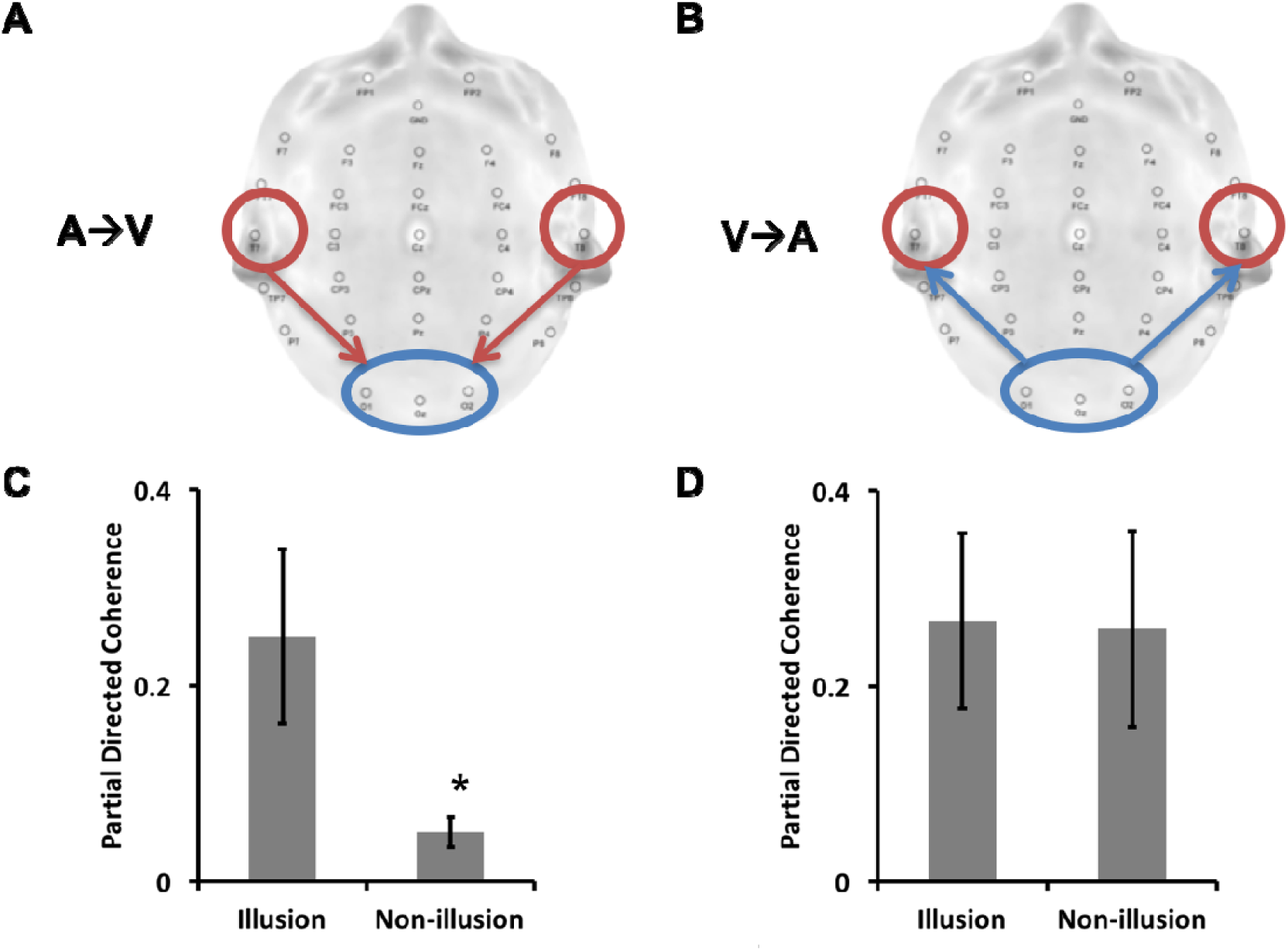
Replication of MEG findings by an independent EEG study, demonstrating higher PDC values from auditory to visual cortices in illusion trials. (A) Partial directed coherence from auditory to visual cortices (A→V), and (B) partial directed coherence from visual to auditory cortices (V→A), in the alpha frequency range (8-12Hz). (C) Partial directed coherence of non-illusion trials decreased significantly compared to that of illusion trials in A→V (\**p*<0.05). (D) They were not different in V→A.

Next, we investigated whether the effective connectivity guided neurofeedback (A→V or V→A) could significantly modulate the sound-induced flash illusion at the behavioral level. We found that after a brief A→V connectivity guided neurofeedback training, participants indeed showed an increased rate of sound-induced visual illusion (Fig. 4). After the A→V neurofeedback training, participants reported significantly higher sound-induced visual illusions in post-training trials with 3 beeps (*t*(26)=8.2 *p*<0.00001) and 4 beeps (*t*(26)=3.0 *p*=.006) (Figs. 4A,B). Further, A→V effective connectivity increased after A→V training (*t*(26)=4.25, *p*=.0002) and decreased after V→A training (*t*(26)=6.66, *p*=0.00001), and this was reflected by an interaction between pre-post and A→V/V→A training, *F*(1,7)=31.6, *p*=0.001. Of note, the number of perceived flashes change after training was marginally correlated with the changes in the A→V cortical PDC values (*R*^2^=0.468, *p*=0.06) (Fig. 4C), yet no such correlation was observed with the changes in the V→A cortical PDC values (*R*^2^=0.247, *p*=0.21) (Fig. 4D).

**Figure 4.**
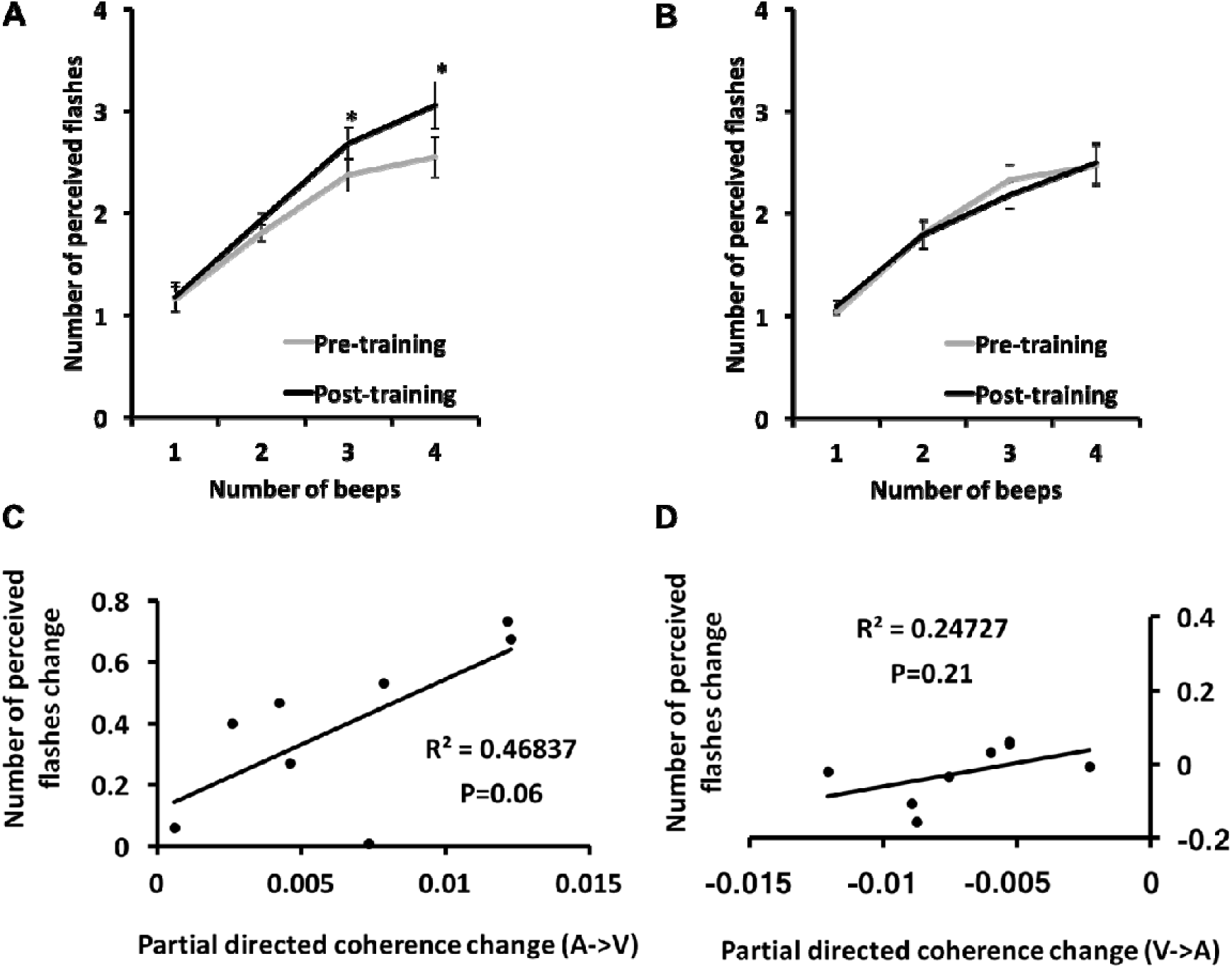
Effective connectivity guided neurofeedback training increases sound-induced visual illusion. (A) Auditory-to-visual training (\**p*<0.05), (B) visual-to-auditory training. Correlations between partial directed coherence change and the number of perceived flashes change in (C) auditory to visual training and in (D) visual to auditory training.

## Discussion

In this study, we demonstrated a robust link between neural dominance and perceptual dominance using sound-induced flash illusion as an experimental paradigm. We showed that effective connectivity from auditory to visual cortices significantly increased in illusion trials compared to non-illusion trials using both EEG and MEG independently. Further, by designing a novel effective connectivity guided neurofeedback protocol, we provided causal evidence that the dominance of the auditory cortex over the visual cortex, but not the other way around, critically influences the reported perceptual dominance of auditory over visual information. Our findings also confirmed the previous findings of increased pre-stimulus auditory and visual connectivity in sound-induced illusion [10]. Our findings also extended the previous findings by providing trial-specific variations, in terms of connectivity between auditory and visual cortices, for identical stimulus configurations, and thereby, establishing a direct link between sensory interactions at the neural level and perceptual outcomes on a trial-by-trial basis. The incorporation of MEG allows a better sensitivity to reveal the connectivity correlates of the sound-induced flash illusion, and the EEG was adopted for the neurofeedback protocol for its practicality and ease of implementation.

Our findings provided evidence for a simple neural mechanism underlying sound-induced visual illusion. Because of the nature of the PDC, which is primarily sensitive to direct functional connections [11], we suggest that the connection from auditory to visual cortices underlies sound-induced flash illusion. However, concluding direct connectivity between two brain regions from EEG/MEG data would remain problematic, so we cannot be certain about the directness of the reported connectivity between the auditory and the visual cortical regions. Further, our sensor selections (i.e. especially the temporal ones) might not reflect activities of purely sensory cortices (i.e. auditory cortex), and the temporal resolution of the frequency domain connectivity, as measured by PDC, should be treated with caution [23]. Nevertheless, we would argue that the ongoing spontaneous interaction of distant cortices, as reported here, could explain the sound-induced visual illusion, and it is possible to alter the qualitative nature of illusory experience by dynamical modulation of the spontaneous effective connectivity between two cortices.

Importantly, we observed a crucial asymmetry between two different directions of neurofeedback training (A→V, V→A). At the neural level, both A→V and V→A training changed the connectivity. However, at the behavioral level, only A→V training led to a significant change. It is consistent with our earlier findings that the sound-induced visual illusion was resistant to feedback training [24]. In other words, the fact that there was only enhancement but no suppression effect might be due to a flooring effect and/or inherent hard connectivity between sensory cortices. Our findings also critically implicate the role of the neural oscillations and effective connectivity, especially in the alpha frequency range [25], subserving multisensory processing [2].

Additionally, we showed that not only can specific regions of the brain be modulated by EEG neurofeedback [22], the connectivity between the regions can also be modulated by the same technique. The connectivity-based neurofeedback is especially useful for establishing a causal relationship between neural activity and behavior. More importantly, this would open ample possible applications whereby training neural connectivity using the feedback technique, we may enhance (or suppress) various mental functions not just limited to multisensory and/or conscious perception.

Summing up, we showed that the spontaneous information flow between sensory cortices as recorded by large scale brain oscillations can be reliably linked with behavioural outcomes, and further, it might be possible to self-regulate this connectivity. These results altogether suggest a more connected and less modular nature of cortical information processing.

## Supplementary Information

**Figure S1.**
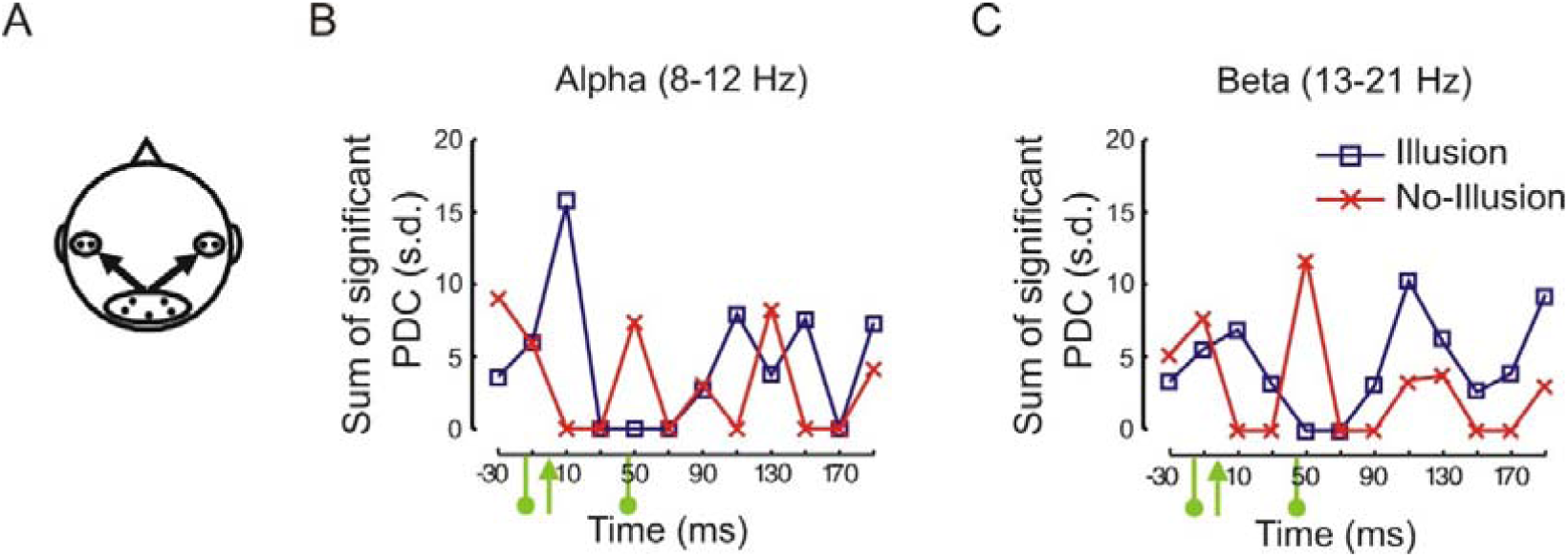
Modulation of auditory cortex by visual cortex. (A) Considered sensors (as in Figure 1B) and direction of information flow. (B)-(C) As in Figure 1C-D, for the causal influence of VC onto AC. As expected (unlike the modulation of the visual cortex by auditory cortex (Figure 1C-D)), no systematically directional influence was observed.

